# Gli2 is necessary for migration of ventral Neural Stem Cells to demyelinated lesions

**DOI:** 10.1101/668418

**Authors:** Daniel Z. Radecki, Heather Messling, James R. Haggerty-Skeans, Jayshree Samanta, James L. Salzer

## Abstract

Enhancing repair of myelin is an important therapeutic goal in many neurological disorders characterized by demyelination. In the healthy adult brain, ventral neural stem cells in the sub-ventricular zone are marked by Gli1 expression and do not generate oligodendrocytes. However, in response to demyelination they migrate to lesions and differentiate into oligodendrocytes. Inhibition of Gli1 further increases their contribution to remyelination. Gli1 and Gli2 are both transcriptional effectors of the Sonic Hedgehog pathway with highly conserved domains but the role of Gli2 in remyelination by ventral neural stem cells is not clear. Here we show that while genetic ablation of Gli1 in the ventral neural stem cells increases remyelination, loss of Gli2 in these cells decreases their migration to the white matter lesion and reduces their differentiation into mature oligodendrocytes. These studies indicate Gli1 and Gli2 have distinct, non-redundant functions in NSCs, including that Gli2 is essential for the enhanced remyelination mediated by Gli1 inhibition. They highlight the importance of designing specific Gli1 inhibitors that do not inhibit Gli2 as a strategy for therapies targeting the Shh pathway.

**Graphical Abstract:** 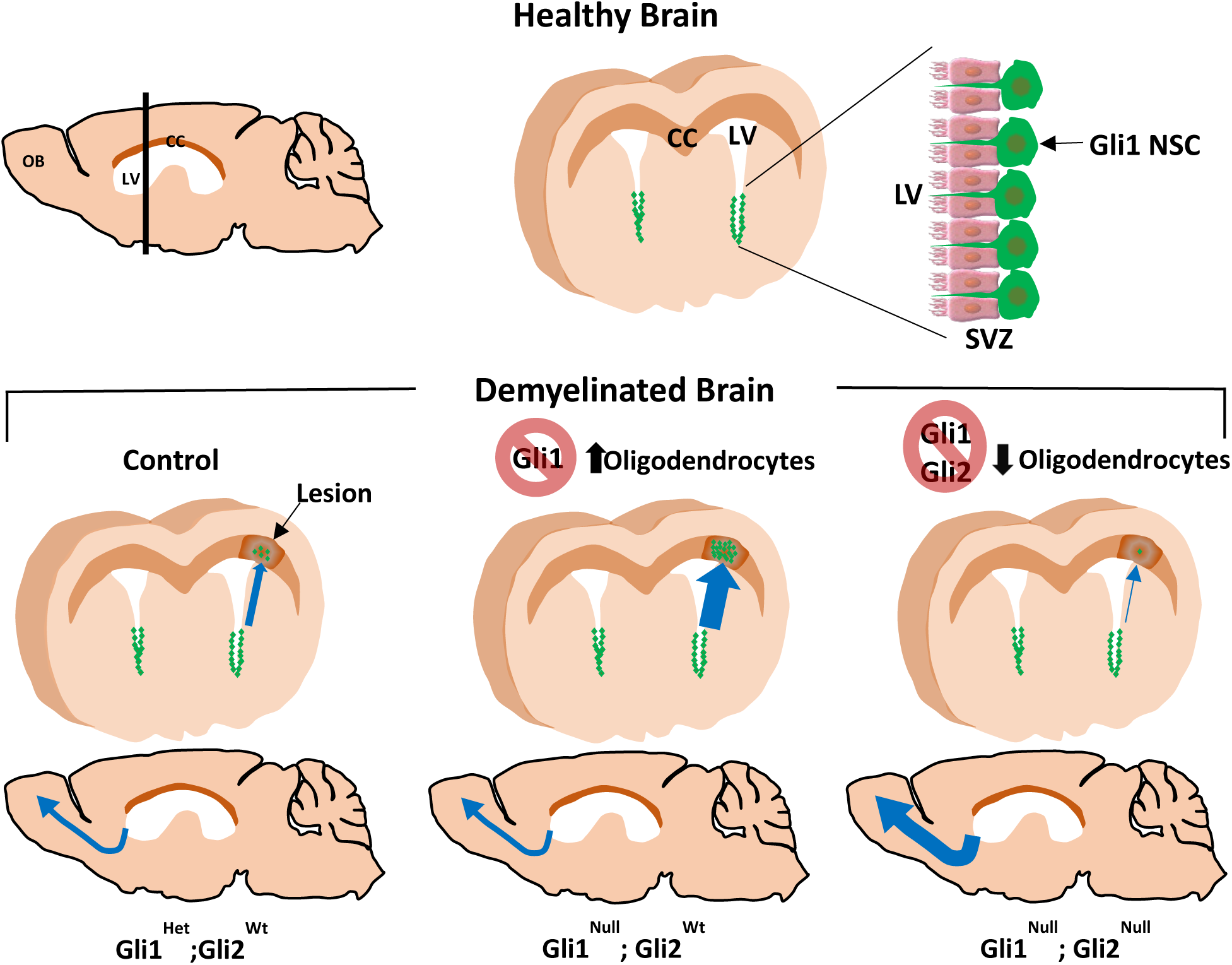

## INTRODUCTION

Neural stem cells (NSC) in the subventricular zone (SVZ) of adult mammalian brains are a heterogeneous population marked by expression of specific transcription factors depending on their embryonic origin (Chaker et al., 2016). A subset of these cells comprising ∼25% of quiescent type B NSCs in the ventral SVZ express Gli1, a transcriptional activator of the Sonic Hedgehog (Shh) pathway (Ahn and Joyner, 2005). Perinatally, these cells are an important source of forebrain oligodendrocytes (Sanchez and Armstrong, 2018). In the adult mouse brain, these ventral NSCs (vNSCs) give rise to neuroblasts, which enter the rostral migratory stream (RMS) and generate interneurons in the olfactory bulb. Outside the SVZ, Gli1 is also expressed by a subset of mature astrocytes; but it is excluded from white matter tracts and not expressed in the oligodendroglial lineage i.e. oligodendrocyte progenitor cells (OPCs) and mature oligodendrocytes (OLs) (Garcia et al., 2010). In contrast, these vNSCs respond to loss of OLs in the white matter by migrating to the demyelinating lesions and differentiating into OLs and astrocytes (Samanta et al., 2015; Sanchez and Armstrong, 2018). Furthermore, the number of cells migrating to the lesions and the proportion of these cells generating remyelinating OLs increases significantly when Gli1 is inactivated genetically or pharmacologically (Samanta et al., 2015).

Gli1 is a zinc-finger transcription factor expressed in response to sustained, high levels of Shh making it a sensitive readout of the pathway. Canonical Shh signaling is mediated by binding of Shh to its receptor Patched, which relieves the inhibition of a G-protein coupled receptor Smo, resulting in activation of the Gli transcription factors. Of the three Gli proteins, Gli1 and Gli2 act as major transcriptional activators (Lipinski et al., 2006). While genetic inactivation of Gli1 does not affect development, knocking out both Gli1 and Gli2 is embryonic lethal suggesting functional redundancy (Bai and Joyner, 2001). In this study, we examined whether Gli2 is necessary for the enhanced remyelination observed by inhibiting Gli1. To overcome the developmental effects, we genetically inactivated Gli2 in the Gli1-pool of vNSCs in adult mice. This ablation impairs the migration to and differentiation of these cells within lesions of the white matter whereas it promotes ongoing migration of vNSCs along the RMS. These results indicate that the normal migration of NSCs along the RMS vs. migration to pathological sites are mechanistically distinct and highlight the non-overlapping activities of Gli1 and Gli2.

## RESULTS

### Gli2 is upregulated in Gli1^NULL^ vNSCs following demyelination

Gli2 is broadly expressed in the NSCs along the entire SVZ suggesting that the increase in remyelination observed on inhibition of Gli1 can potentially result from non-canonical activation of Gli2 (Petrova et al., 2013). Hence, we first compared the expression of Gli2 in the SVZ of brains expressing Gli1 (Gli1^HET^) vs those lacking Gli1 (Gli1^NULL^) by qRT-PCR after inducing demyelination with Cuprizone diet (Matsushima and Morell, 2001). There was a trend towards increased Gli2 expression with demyelination in the Gli1-Hets, which was significant in the Gli1-Nulls. In particular, there was a significant increase (2.3±0.37 fold) in the levels of Gli2 mRNA in the Gli1^NULL^ SVZ at peak demyelination (6wks cuprizone diet) as compared to the healthy SVZ (regular diet) suggesting a positive role in remyelination (Fig. 1A,C). To examine the expression of Gli2 further, we crossed the Gli1^HET^ and Gli1^NULL^ mice with Gli2^nLacZ^ mice which express nuclear LacZ in all Gli2-expressing cells (Bai and Joyner, 2001). Using the RCE reporter to fate-map Gli1 vNSCs (Samanta et al., 2015), we observed co-expression of Gli2 in a subset of fate-mapped Gli1^HET^ and Gli1^NULL^ NSCs in the SVZ of healthy as well as demyelinated brains (Fig.1A,B). Consistent with our prior results showing higher proliferation in Gli1^NULL^ vNSCs vs Gli1^HET^ vNSCs upon demyelination (Samanta et al., 2015), we found a qualitative increase in the number of Gli1^NULL^ vNSCs co-expressing Gli2. These data suggest that loss of Gli1 increases the expression of Gli2, which may influence remyelination by the vNSCs.

**Figure 1.**
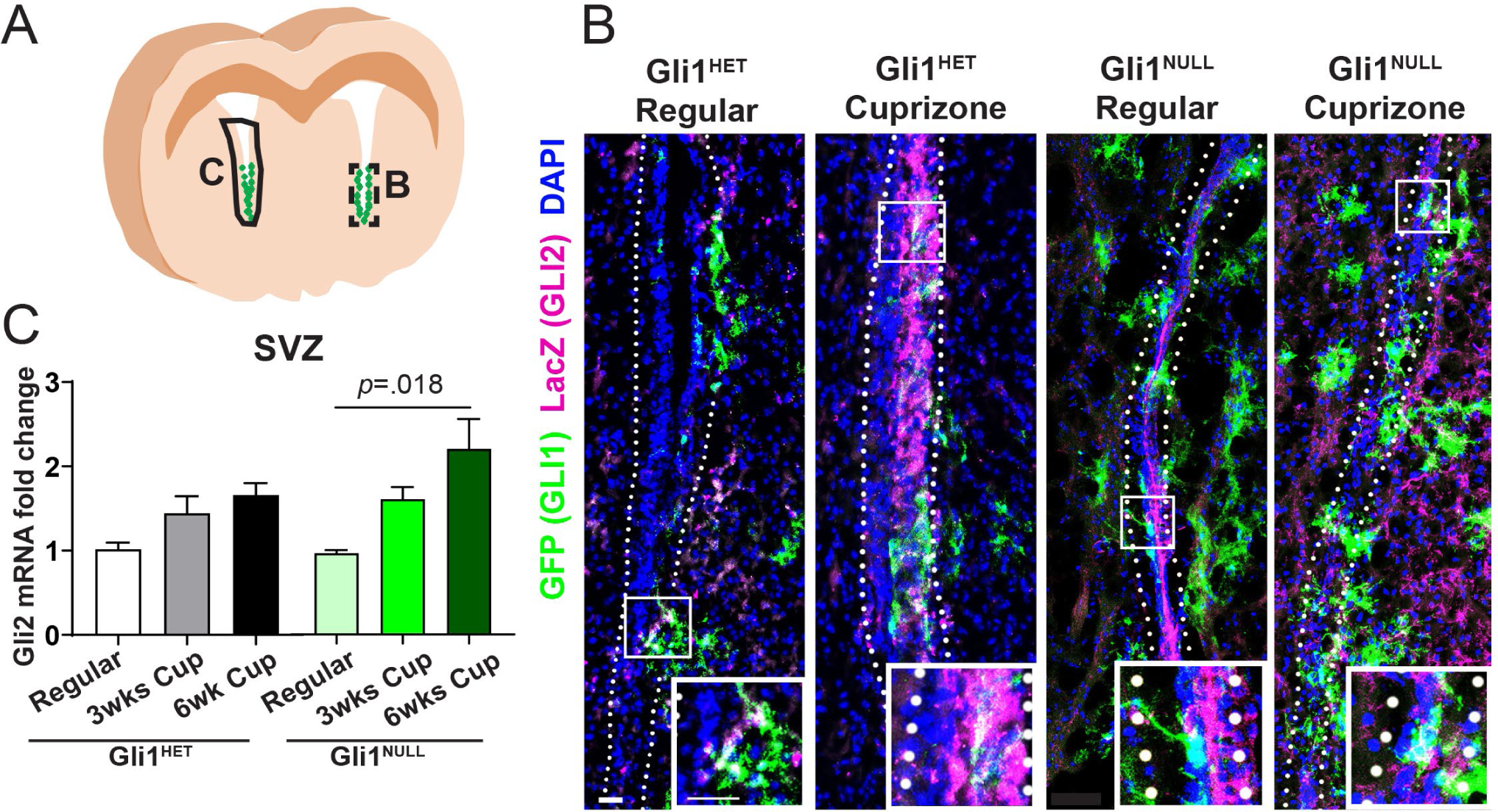
Gli2 expression increases in the Gli1^NULL^ SVZ on demyelination. A) Outline of coronal brain section showing the tissue harvested for immunofluorescence (B) and qRT-PCR (C). B) Immunofluorescence of the SVZ (dotted lines) shows expression of GFP (green) in Gli1 fate-mapped cells and LacZ (magenta) in Gli2+ cells in Gli1 ^HET^ and Gli1^NULL^ mice on 6 weeks of regular or cuprizone diets. Scale = 50µm. C) Gli2 mRNA expression is increased in the Gli1^NULL^ SVZ at 6 weeks of demyelination. 1-way ANOVAs with Tukey’s post-hoc t-tests; data presented as mean ± SEM; n=3; subventricular zone (SVZ), Cuprizone diet (Cup)

### Loss of Gli2 decreases the migration of vNSCs to white matter lesions

To determine the role of Gli2 in remyelination, we examined the effects of different amounts of Gli2 expression on vNSC remyelination. In particular, we conditionally inactivated Gli2 in the Gli1-Het and Null vNSCs and then analyzed their fate in the healthy brain (mice on a regular diet) or at 2 weeks of recovery after Cuprizone-induced demyelination (Fig. 2A,B). As we reported previously (Samanta et al., 2015), there were no fate mapped cells in the healthy corpus callosum (CC) and many more Gli1-Null vs. Het NSCs in the CC following demyelination. We further observed a trend towards a gene dosage dependent decrease in the number of fate-mapped cells in the CC of Gli1^HET^ mice with loss of one or both alleles of Gli2 compared to Gli2^WT^ mice. In contrast, the Gli1^NULL^ mice heterozygous for Gli2 showed a significant 83.7±2.4% reduction of fate-mapped vNSCs, while complete loss of Gli2 resulted in an 89.8±1.9% reduction in the GFP-labeled cells entering the CC compared to the Gli2^WT^ mice (Fig. 2A, B). Thus, haploinsufficiency of Gli2 in the Gli1^NULL^ vNSCs is sufficient to reverse their enhanced recruitment into the CC upon demyelination.

**Figure 2.**
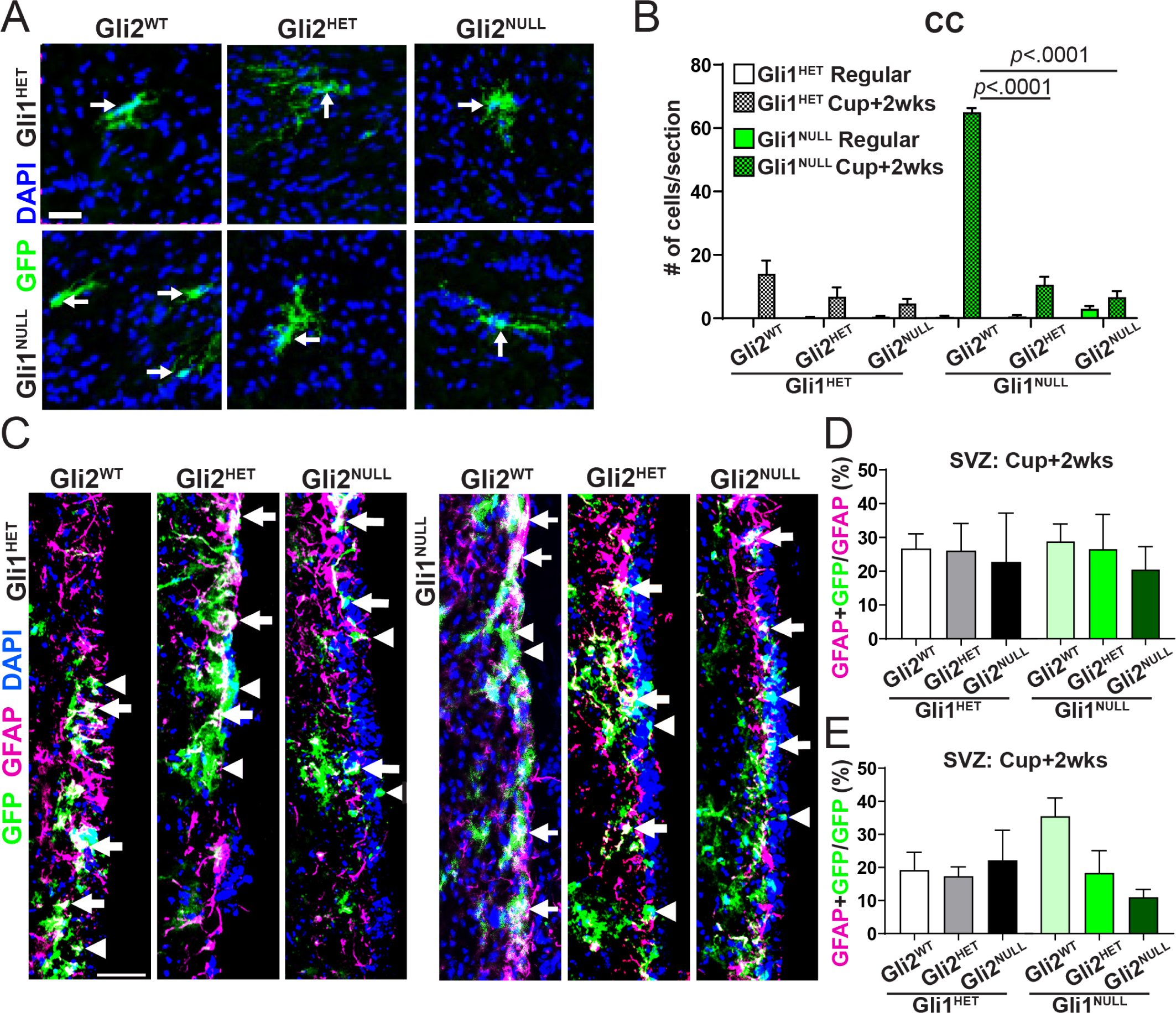
Ablation of Gli2 decreases migration of Gli1^NULL^ vNSCs to the white matter lesion. A) Immunofluorescence for fate-mapped Gli1^HET^ (top) and Gli1^NULL^ (bottom) vNSCs in the CC at 2 weeks of remyelination (arrows). Scale = 50µm. B) Quantification of the total number of fate-mapped Gli1^HET^ (white) and Gli1^NULL^ (green) vNSCs in the CC shows a reduction with loss of Gli2 in Gli1^NULL^ mice. 2-way ANOVAs with Tukey’s post-hoc t-tests within Gli1 groups; mean ± SEM; n=5. C) Immunofluorescence for GFAP (magenta) labelling quiescent NSCs and GFP (green) in fate-mapped Gli1^HET^ (left) or Gli1^NULL^ (right) vNSCs in the SVZ. Gli1+/GFAP+ NSCs (arrows); Gli1+/GFAP-cells (arrowheads). Scale bar = 50µm. D) The proportion of GFAP+ NSCs in the SVZ that were fate-mapped Gli1 vNSCs co-expressing GFAP was unchanged by loss of Gli2 at 2 weeks of remyelination. E) The proportion of fate-mapped Gli1 vNSCs that co-expressed GFAP was unaltered by loss of Gli2 at 2 weeks of remyelination. For D and E: multiple t-tests within Gli1 groups and comparing all groups to Gli1^HET^;Gli2^WT^; data presented as mean ± SEM; n=3. Ventral neural stem cells (vNSCs), corpus callosum (CC), subventricular zone (SVZ), Cuprizone diet (Cup)

Although loss of Gli2 does not alter neurogenesis from vNSCs in the healthy SVZ (Petrova et al., 2013), its role in maintaining quiescence of these cells upon demyelination is unknown. Hence, we quantified GFP-labeled cells in the SVZ that co-express GFAP, a NSC marker, and did not detect any difference in the relative proportion of vNSCs at 2 weeks of recovery from Cuprizone in Gli1^HET^ and Gli1^NULL^ mice (Fig. 2C,D). Reciprocally, there was no change in the fraction of the Gli1-pool that were quiescent vNSCs in the SVZ (Fig. 2E). These results indicate that loss of Gli2 does not alter the vNSC population in the SVZ of Gli1^HET^ and Gli1^NULL^ mice following demyelination.

### Loss of Gli2 increases migration of NSCs to the olfactory bulb on demyelination

Taken together, our findings indicate that Gli2 is necessary for migration of vNSCs to the white matter lesions for remyelination. They raise the possibility that Gli2 is required for vNSCs migration more broadly. To address this, we examined migration of the Gli1 pool of vNSCs from the SVZ to the olfactory bulb (OB) in parallel during remyelination. vNSCs are known to produce deeper layer granule interneurons in the OB whereas dorsal NSCs that do not express Gli1, generate interneurons of the superficial granule layer in the OB (Ihrie et al., 2011; Young et al., 2007). Loss of Gli2 from all the NSCs in the healthy SVZ results in a decrease in the superficial granule layer interneurons without affecting the deeper layers suggesting that Gli2 is not essential for vNSC migration (Petrova et al., 2013). However, the role of Gli2 in the migration of vNSCs that lack Gli1 to the OB and upon demyelination is not clear.

In general, we observed similar numbers of *Gli1*^*CreERT2*^ fate-mapped cells in the OBs of Gli1^HET^ and Gli1^NULL^ mice on regular diet (data not shown). We next quantified the numbers of such GFP-labeled cells in the OB of Gli1^HET^ and Gli1^NULL^ brains with varying levels of Gli2 expression at 2 weeks of recovery from demyelination. There was a significant increase in the number of fate-mapped cells in the Gli1^NULL^ OBs with loss of one or both alleles of Gli2 (i.e. 235.5±38 cells/mm^2^ in Gli2^HET^ and 222.8±23.4 cells/mm^2^ in Gli2^NULL^) compared to Gli2^WT^ controls (93.4±20.8 cells/mm^2^; Fig. 3A,B). However, the position and morphology of labeled cells within the OB was not affected by loss of Gli2. Cells with a neuronal morphology were located in the deeper granule cell layers and those with an astrocytic morphology were predominantly in the external plexiform layer of the OB (Fig. 3A,a’). To determine if there were any effects on the number of vNSCs, we also quantified all the fate-mapped cells that had migrated out of the SVZ and found that loss of Gli2 did not alter the combined numbers of GFP-labeled cells in the CC and OB (Fig.3C). Together these data show that in the demyelinated brain, inhibition of both Gli1 and Gli2 in vNSCs promotes their migration towards the OB and away from the lesion.

**Figure 3.**
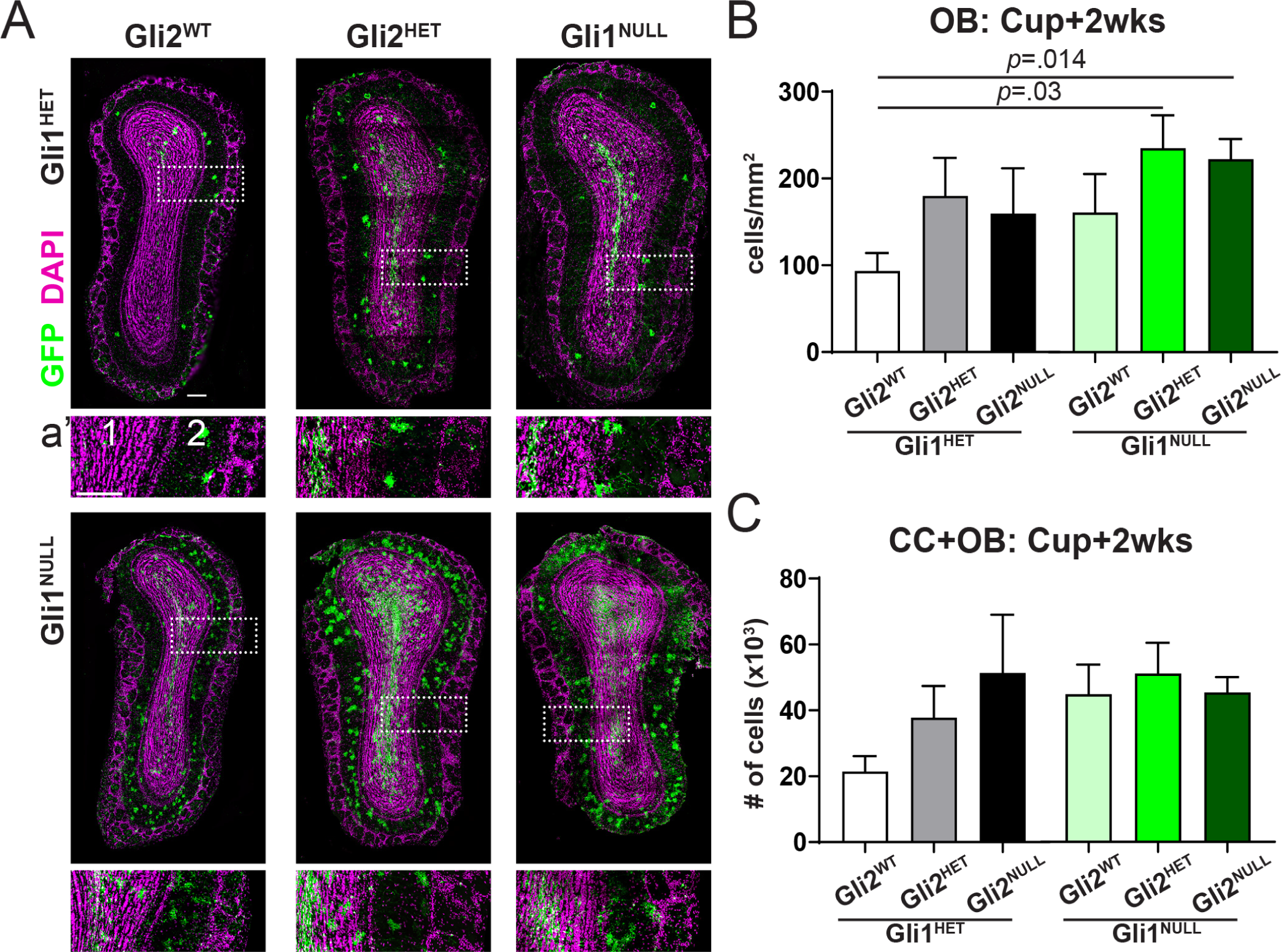
Ablation of Gli2 increases migration of Gli1^NULL^ vNSCs to the olfactory bulb. A) Immunofluorescence for fate-mapped Gli1^HET^ (top) and Gli1^NULL^ (bottom) vNSCs (green) in the OB at 2 weeks of remyelination shows an increase in Gli1^NULL^;Gli2^HET^ and Gli1^NULL^;Gli2^NULL^ mice compared to Gli1^HET^;Gli2^WT^ controls. **a’)** Enlarged portion of the OB (dotted rectangle) showing (1) granule and (2) external plexiform layers. B) Quantification of the fate-mapped Gli1^HET^ and Gli1^NULL^ vNSCs in the OB shows an increase in Gli1^NULL^;Gli2^HET^ and Gli1^NULL^;Gli2^NULL^ mice compared to Gli1^HET^;Gli2^WT^ controls. C) Quantification of the fate-mapped Gli1^HET^ and Gli1^NULL^ cells in the combined areas (CC and OB), shows similar numbers at 2 weeks of remyelination. Multiple t-tests within Gli1 groups and comparing all groups to Gli1^HET^;Gli2^WT^ controls; data presented as mean ± SEM; n=3; Scale = 50µm; Ventral neural stem cells (vNSCs), olfactory bulb (OB), corpus callosum (CC), subventricular zone (SVZ), Cuprizone diet (Cup)

### Gli2 promotes the differentiation of Gli1-null vNSCs into OLs in the lesion

In the healthy brain, Gli2 is expressed by all NSCs in the SVZ, and by mature astrocytes, but not by cells in the OL lineage (Garcia et al., 2010). Accordingly, we examined if Gli2 affects the generation of OLs from vNSCs in response to demyelination in the CC. In the Gli1^HET^ CC, we observed a significant reduction in the generation of astrocytes whereas the proportion of OLs and OPCs was not altered upon ablation of Gli2 (Fig. 4A,C,D). In contrast, we observed a ∼50% reduction in the generation of mature OLs (mature OL marker CC1 and myelin marker MBP), when Gli2 was lost in the Gli1^NULL^ mice. Specifically, the proportion of mature OLs was reduced from 81.2±2.2% of fate-mapped cells in the Gli2^WT^ brains to 24.9±11.2% in Gli2^HET^ brains and 33.2±12.3% in Gli2^NULL^ brains (Fig. 4A,B). However, there were no significant differences in the proportion of astrocytes in the Gli1^NULL^ mice (Fig. 4C). Consistent with our previous study, we also found that at 2 weeks of recovery from cuprizone diet, a considerable number of fate-mapped vNSCs did not express any markers of differentiation in the lesion site (Samanta et al., 2015). Overall, these data indicate that Gli2 is necessary for the enhanced generation of mature OLs in Gli1^NULL^ mice during remyelination.

**Figure 4.**
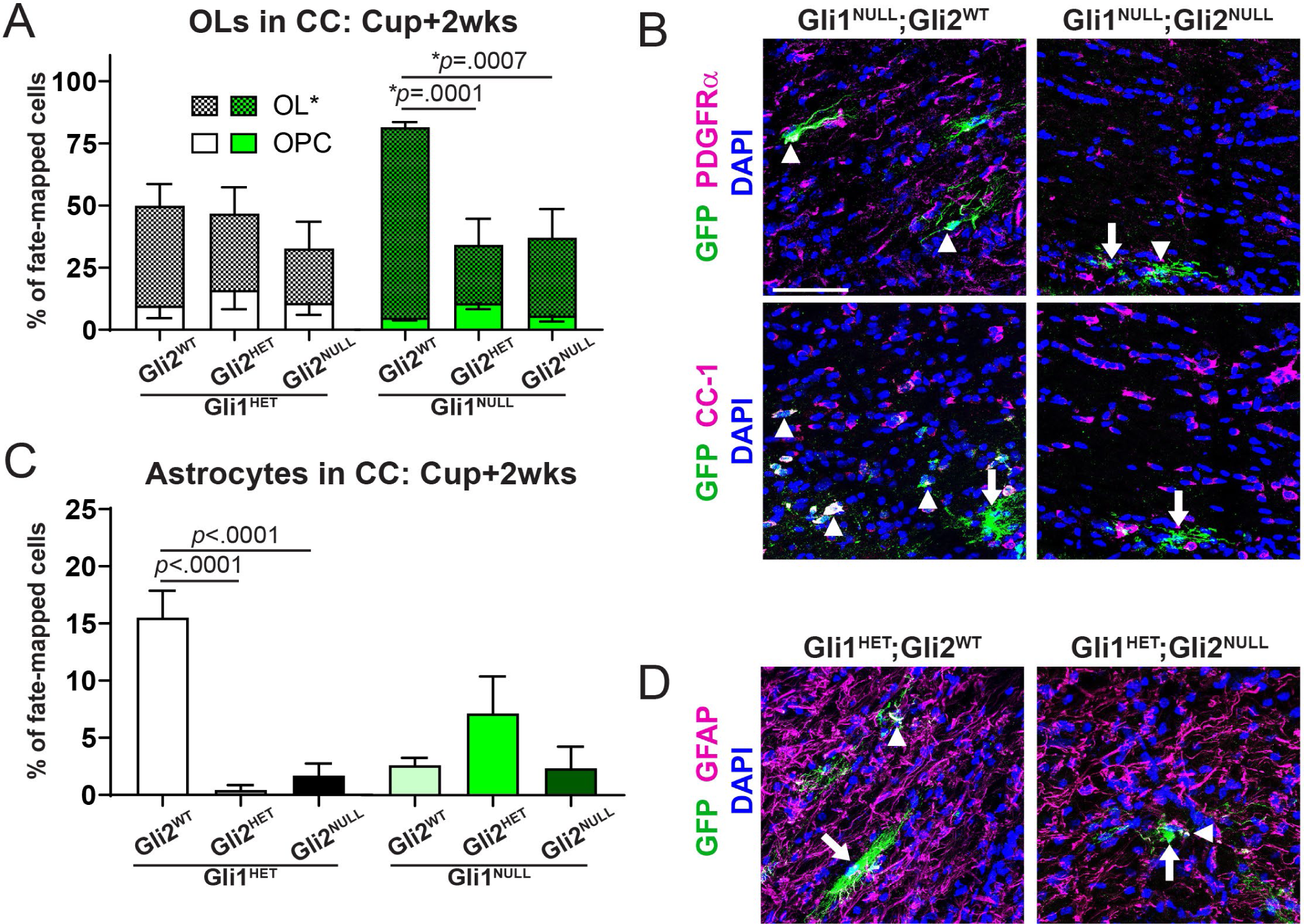
Loss of Gli2 alters the differentiation of vNSCs in the corpus callosum. A) Quantification of the percentage of fate-mapped Gli1^HET^ (white) and Gli1^NULL^ (green) vNSCs that co-express markers of OPC (PDGFRα) and OL (CC1+MBP) shows a decrease in OLs in the Gli1^NULL^ CC with loss of one or both copies of Gli2. B) Immunofluorescence for fate-mapped Gli1^NULL^ vNSCs (green) co-expressing markers (magenta) of OPC(PDGFRα) and OL (CC1) shows a decrease in OLs upon loss of Gli2. C) Quantification of the percentage of fate-mapped Gli1^HET^ (white) and Gli1^NULL^ (green) vNSCs that co-express the astrocyte marker (GFAP) shows a decrease in the Gli1^HET^ CC with loss of one or both copies of Gli2. D) Immunofluorescence for fate-mapped Gli1^HET^ vNSCs (green) co-expressing the astrocytic marker GFAP (magenta) shows a decrease upon loss of Gli2. 1-way ANOVA with Tukey’s post-hoc t-tests within Gli1 genotypes; all data presented as mean ± SEM; n=3-5 for all; scale bar = 50µm. Fate-mapped cells (arrows) and co-expressing cells (arrowheads). Ventral neural stem cells (vNSCs), corpus callosum (CC), oligodendrocyte progenitor cells (OPC), oligodendrocyte (OL), Cuprizone diet (Cup)

## DISCUSSION

This study shows that Gli2 plays an essential role in promoting remyelination by vNSCs, particularly in the absence of Gli1. We observed an increase in Gli2 expression following demyelination in Gli1-deficient SVZ. When we ablated Gli2 in Gli1 vNSCs, the resulting progeny preferentially migrated to the OB instead of the demyelinated CC. This altered migration pattern suggests that Gli2 expression may normally divert vNSCs away from the RMS and the OB.

We had previously shown that Gli1^NULL^ vNSCs exhibit increased OL generation and decreased astrocyte differentiation in the demyelinated CC compared to Gli1^HET^ vNSCs (Samanta et al., 2015). Data in this study indicate Gli2 is essential for differentiation of Gli1^HET^ vNSCs into astrocytes – in its absence, the generation of astrocytes is reduced to the low levels exhibited by Gli1^NULL^ vNSCs. Loss of Gli2 in the Gli1^NULL^ vNSCs does not further reduce their differentiation into astrocytes from an already low baseline. Similarly, complete loss of Shh signaling in the healthy brain leading to loss of both Gli1 and Gli2 activities, results in an activated phenotype without a change in the number of astrocytes (Garcia et al., 2010). Our data further shows that loss of Gli2 does not reduce oligodendrogenesis by Gli1^NULL^ vNSCs beyond that seen in Gli1^HET^ vNSCs upon demyelination.

Taken together, our results indicate Gli2 is necessary for the enhanced migration and oligodendrogenesis in the white matter lesions when Gli1 is ablated. These data underscore that Gli1 and Gli2 have distinct functions in the fate of vNSCs in response to demyelination, i.e. loss of Gli1 promotes their efficacy in remyelination whereas loss of Gli2 inhibits their efficacy. These latter findings are consistent with the distinct targets these two related transcription factors regulate (Ali et al., 2019). These results further suggest that therapeutic strategies that target the transcriptional effectors of Shh (Lauth et al., 2007) could have very different outcomes depending on the relative specificity for Gli1 vs. Gli2.

## EXPERIMENTAL PROCEDURES

### Fate mapping and demyelination

All animals were used and maintained according to protocols approved by the University of Wisconsin IACUC. These mouse lines were obtained from Jackson labs: Gli1^CreERT2^ (Jax# 007913), Rosa-CAG-EGFP (RCE) (Jax# 032037); Gli2^nLacZ^ (Jax# 007922) and Gli2^Flox^ (Jax#007926). The genotypes for the mouse lines are as follows: Gli1^HET^-Gli1^CreERT2/+^;RCE, Gli1^NULL^-Gli1^CreERT2/CreERT2^;RCE, Gli2^HET^-Gli2^Flox/+^ or Gli2^LacZ/+^ and Gli2^NULL^-Gli2^Flox/Flox^. All the mice were maintained on C57Bl/6 background. 10 week old mice were administered 5 mg tamoxifen (Sigma) in corn oil on alternate days for a total of four intraperitoneal injections. No labelling was seen in the absence of tamoxifen administration. A week later, demyelination was induced by feeding 0.2% cuprizone in the chow for 6 weeks following which the diet was returned to normal chow for recovery from demyelination.

### Immunostaining

Mice were perfused with 4%PFA, and 20 µm coronal cryosections were processed for immunofluorescence with rabbit or chicken anti-GFP (1:1,000, Invitrogen) and one of the following antibodies: rat anti-PDGFRα (1:200, BD Biosciences); mouse anti-CC1 (1:400, Millipore), anti-GFAP (1:400, Sigma), anti-MBP (1:500, Millipore) and anti-LacZ (1:2,000, Sigma). Secondary antibodies were donkey or goat anti-species conjugated with Alexafluors (1:1,000, Molecular probes). Nuclei were counterstained with Hoechst 33258 (1:5,000, Invitrogen). Epifluorescent images were obtained as Z-stacks of 5µm optical sections using a Keyence BZ-X700 microscope at 20x magnification. For quantification of CC1, PDGFRα, GFAP, MBP and OB GFP immunostaining, ImageJ cell counting feature was used. Keyence BZ-X Analyzer Hybrid Cell Count tool was used to analyze the SVZ. Publication images were obtained as Z-stacks of 1μm optical sections using a confocal laser-scanning microscope (Leica TCS SP8, LasX software) and processed using Adobe Photoshop. The investigators were blinded to allocation during experiments and outcome assessment.

### qRT-PCR

mRNA was extracted from the SVZ of three mice in each group using RNeasy kit (Qiagen) and reverse-transcribed to complementary DNA using iScript cDNA Synthesis Kit (BioRad). ITaq Universal SYBR Green Supermix (BioRad) was used to perform qPCRs in a Biorad CFX Connect thermal cycler. Primers used were Gli2 (forward, 5’-AGA GAC AGC AGA AGC TAT GCC CAA-3’; reverse, 5’-TGG GCA GCC TCC ATT CTG TTC ATA-3’) and GAPDH (forward, 5’-GGT GTG AAC GGA TTT GGC CGT ATT G-3’; reverse, 5’-CCG TTG AAT TTG CCG TGA GTG GAG T-3’). The 2^-ΔΔCt^ method was used to analyze the relative gene expression, each Gli1 genotype was normalized to regular diet.

### Statistical Analysis

All results were validated by at least three independent experiments. At least 5 sections per mouse were analyzed and data from 3–5 mice were combined to determine the average and standard deviation and SEM. The quantitative data were expressed as mean ± SEM. Statistical analysis was performed using Student’s t test, 1-way ANOVA, or 2-way ANOVA with Tukey’s post hoc test. Differences were considered statistically significant at p < 0.05.

## AUTHOR CONTRIBUTIONS

D.Z.R. and H.M. performed research, analyzed and interpreted results and edited the manuscript; J.R.H. performed preliminary cell counts; J.S. designed and performed research, analyzed and interpreted results, and wrote the manuscript; J.L.S. reviewed data and edited the manuscript.

## ACKNOWLEDGEMENTS

A patent on the method of targeting Gli1 as a strategy to promote remyelination has been awarded, with J. L. Salzer, J. Samanta and G. Fishell listed as co-inventors. GliXogen Therapeutics Ltd. has licensed the technology from New York University. This research was supported by grants to J.L.S. from the New York State Department of Health Stem Cell Board and to J.S. from Nancy and Jean-Pierre Boespflug Foundation.

